# Evidence of the absence of Human African Trypanosomiasis in northern Uganda: analyses of cattle, pigs and tsetse flies for the presence of Trypanosoma brucei gambiense

**DOI:** 10.1101/753020

**Authors:** Lucas J. Cunningham, Jessica K. Lingley, Iñaki Tirados, Johan Esterhuizen, Mercy A. Opiyo, Clement T. N. Mangwiro, Mike J. Lehane, Stephen J. Torr

## Abstract

**Background:** Large-scale control of sleeping sickness has led to a decline in the number of cases of Gambian human African trypanosomiasis (g-HAT) to <2000/year. However, achieving complete and lasting interruption of transmission may be difficult because animals may act as reservoir hosts for *T. b. gambiense*. Our study aims to update our understanding of *T. b. gambiense* in local vectors and domestic animals of N.W. Uganda.

**Methods:** We collected blood from 2896 cattle and 400 pigs and In addition, 6664 tsetse underwent microscopical examination for the presence of trypanosomes. *Trypanosoma* species were identified in tsetse from a subsample of 2184 using PCR. Primers specific for *T. brucei* s.l. and for *T. brucei* sub-species were used to screen cattle, pig and tsetse samples.

**Results:** In total, 39/2,088 (1.9%; 95% CI=1.9-2.5) cattle, 25/400 (6.3%; 95% CI=4.1-9.1) pigs and 40/2,184 (1.8%; 95% CI=1.3-2.5) tsetse, were positive for *T. brucei* s.l.. Of these samples 24 cattle (61.5%), 15 pig (60%) and 25 tsetse (62.5%) samples had sufficient DNA to be screened using the *T. brucei* sub-species PCR. Further analysis found no cattle or pigs positive for *T. b. gambiense*, however, 17/40 of the tsetse samples produced a band suggestive of *T. b. gambiense*. When three of these 17 PCR products were sequenced the sequences were markedly different to *T. b. gambiense*, indicating that these flies were not infected with *T. b. gambiense*.

**Conclusion:** The absence of *T. b. gambiense* in cattle, pigs and tsetse accords with the low prevalence of g-HAT in the human population. We found no evidence that livestock are acting as reservoir hosts. However, this study highlights the limitations of current methods of detecting and identifying *T. b. gambiense* which relies on a single copy-gene to discriminate between the different sub-species of *T. brucei* s.l.

**Author Summary:** The decline of annual cases of West-African sleeping sickness in Uganda raises the prospect that elimination of the disease is achievable for the country. However, with the decrease in incidence and the likely subsequent change in priorities there is a need to confirm that the disease is truly eliminated. One unanswered question is the role that domestic animals play in maintaining transmission of the disease. The potential of cryptic-animal reservoirs is a serious threat to successful and sustained elimination of the disease. It is with the intent of resolving this question that we have carried out this study whereby we examined 2088 cattle, 400 pigs and 2184 tsetse for *Trypanosoma brucei gambiense*, the parasite responsible for the disease. Our study found *T. brucei* s.l. in local cattle, pigs and tsetse flies, with their respective prevalences as follows, 1.9%, 6.3% and 1.8%. Further analysis to establish identity of these positives to the sub-species level found that no cattle, pigs or tsetse were carrying the pathogen responsible for Gambian sleeping sickness. Our work highlights the difficulty of establishing the absence of a disease, especially in an extremely low endemic setting, and the limitations of some of the most commonly used methods.

## Introduction

The term “human African trypanosomiasis” (HAT) is used to describe two diseases that are clinically, geographically and parasitological distinct. The majority of HAT cases (98%) occur in West and Central Africa and are referred to as West African sleeping sickness or Gambian HAT (g-HAT) indicating the geographical range of the disease and the protozoan parasites responsible, *Trypanosoma brucei gambiense*. Similarly, East African sleeping sickness or Rhodesian HAT (r-HAT) results from an infection caused by *T. b. rhodesiense*. While *T. b. rhodesiense* has long been known to have a primarily zoonotic lifecycle, *T. b. gambiense* is considered to be largely anthropophilic with the parasites largely circulating between tsetse and humans only. *T. b. gambiense* has been identified in domestic animals such as pigs, sheep and goats (1, 2) as well as in a number of wild animals (3, 4). Similarly, a wide range of animals have been experimentally infected with *T. b. gambiense* some of which were shown to be infective to tsetse. These observations suggest that it may be possible for animals to act as reservoirs hosts for *T. b. gambiense* (5–7) and play a role in transmission. Another study that supports the possibility of cryptic animal reservoirs are the reports of tsetse infected with *T. b. gambiense* caught in areas without cases of g-HAT (8). The unresolved question, of a zoonotic host, in the life history of *T. b. gambiense* has significant consequences if elimination by 2030 is to be achieved (9). The importance of a successful elimination campaign that does not result in low prevalence pockets of transmission is evident when one considers the history of HAT. Since the turn of the 20^th^ century there have been three major outbreaks of sleeping sickness resulting in hundreds of thousands of deaths. Crucially the third outbreak occurred after intense control efforts had reduced the number of HAT cases to near-elimination levels (10). The threat of resurgence will always be present and require continued pressure to keep HAT in check unless it is truly eradicated.

Despite the well documented reports of animals infected with *T. b. gambiense*, and the evidence that tsetse can become infected through animal hosts, it is not known if zoonotic cases of *T. b. gambiense* act as cryptic reservoirs that play a role in sustaining transmission of gHAT. Modelling studies (11–13) have shown that the success or failure of eliminating sleeping sickness depends on a number of parameters, one of which is the existence of a cryptic-animal reservoir. The presence of an animal reservoir can also change the importance of the other parameters such as the importance of human migration to an area with a high tsetse biting rate in the context of heterogenous biting (12).

A limiting factor to analysing the role and importance of non-human hosts is the type of diagnostic method used to detect the presence of trypanosomes. Classically, microscopic detection of parasites in blood of a human is regarded as evidence of infection. However, for animal hosts this method is unable to distinguish human-infective *T. b. gambiense* from animal-infective *T. b. brucei* (14). Molecular methods can reliably distinguish the different trypanosome species, with a high degree of sensitivity and specificity (15). However, differentiation of the *T. brucei* sub-species, although possible, has limited sensitivity, as single copy regions of the genome must be targeted (16). Ideally samples positive for *T. brucei* s.l. will need to be assessed to verify there is enough DNA present to undergo the less sensitive sub-species PCR assay (17). To date the application of these molecular methods has not been fully applied to the N.W. of Uganda, although an animal survey was conducted from 2004-2008, this study did not validate the samples suitability of single copy gene amplification (18). It is likely that a potion of those samples identified as being positive for *T. brucei* s.l. lack sufficient DNA to undergo the sub-species-specific detection assay.

An alternative to molecular methods are serological techniques including the card agglutination test for trypanosomiasis (CATT) and the trypanolysis (TL) test, however the CATT can produce false positives due to malaria and transient trypanosome infections (19, 20).The sensitivity of the two methods also varies between geographical locations (21). The unreliability of these methods can vary across different geographical areas due to the heterogenic distribution of the markers in wild trypanosome populations, making their reliability variable (22).

### Aim

Here we use currently available molecular assays to determine the presence or absence of *T. b. gambiense* in NW Uganda by screening the vector population and two potential, animal-reservoir populations, cattle and pigs, in a large-scale xenomonitoring campaign, using molecular methods to first identify cases of *T. brucei* s.l. and subsequently the sub-species of *T. brucei* with PCR assays.

## Methods

### Study site

The North West of Uganda has nine districts, Nebbi, Arua, Koboko, Yumbe, Moyo, Adjumani, Maracha, Amuru and Gulu, of which Arua, Koboko, Yumbe, Moyo, Maracha, Amuru and Adjumani have historic sleeping sickness foci (23). Records from 1905-1920 show deaths from HAT in the West Nile region to be 1-100 per 10,000 (24). Recent surveys from 2000-2015 show that this area of Uganda still has foci of disease, (25) recently there has been a decrease in new cases of HAT being reported and only three new cases in 2016. In 2018, 10,000 people from 28 villages across four districts (Arua, Maracha, Koboko and Yumbe) of N.W. Uganda were screened using the card agglutination test for trypanosomiasis (CATT), with any positives being followed up by CTC microscopy and trypanolysis. Out of the 10,000 individuals screened none were found to have a current *T. b. gambiense* infection, although three participnats did test positive for having had *T. b. gambiense* in the past (26). In this study tsetse were caught in the district of Koboko and screened for trypanosomes and in both Koboko and Arua cattle were sampled and screened for trypanosomes. No sampling for tsetse was carried out in Arua due to a vector control programme being carried out in this district (27). Pigs were sampled from Arua but not from Koboko as there are few pigs there due to it being predominantly Muslim and hence domestic pigs are scant.

### Tsetse catches

A total of 12,152 tsetse flies were caught along the Kochi River in the district of Koboko over a period of 16 months from April 2013 to July 2014. To catch tsetse, pyramidal traps (28) were deployed at four locations (383200N-283715E, 381611N-287545E, 383674N-280855E, 383550N-283841E). The traps were located <1m from the river and tsetse were collected twice a day, at ^∼^07:30 h and ^∼^15:30 h. Tsetse were transported from the trap sites to the field laboratory in cool boxes containing a damp towel to reduce heat and maintain humidity to reduce mortality of tsetse prior to dissection.

Of the tsetse caught, 6,664 were dissected at the field laboratory (333842N-269418E) to screen for trypanosomes in their mouthparts, salivary glands and midguts (29). Following their dissection, the three tissue types (mouthparts, salivary glands and midguts) were screened for trypanosomes at x400 using a compound-microscope with a dark-field filter. Both negative and positive tissues were then preserved, individually, in 100% ethanol, on a 96 well plate, for further molecular analysis. Of the 6,664-tsetse preserved in this manner 2,184 were processed using the molecular methods described below. The sub-sample of tsetse was selected based on the season they were caught, with samples from the wet and dry season of Septemeber2013-February 2014 being chosen.

### Tsetse DNA extraction

After transportation to the Liverpool School of Tropical Medicine, at room temperature, samples were stored at 4°C until undergoing DNA extraction, after which the samples were stored, in a separate fridge, at 4°C. Each individually-preserved tsetse tissue underwent DNA extraction, previously described in Cunningham *etal.* (30). Briefly, ethanol was evaporated by incubating the samples at 56°C for 3 hours and to the sample 135μl of extraction buffer was added (1% Proteinase K, 5% TE Chelex suspension). Finally, the samples underwent incubation at 56°C for one hour followed by incubation at 93°C for 30 minutes to halt the enzymatic activity of the proteinase K.

### Cattle and Pig sampling

From August 2011 to December 2013, 2,896 cattle blood samples were collected across seven sites in N.W. Uganda, as part of an impact assessment study following the deployment of tiny targets to control the local tsetse population (27). Of the 2,896 cattle samples taken 2088 were screened for *T. brucei* s.l. using the FIND-TBR primers (27). Alongside the cattle sampling, in August 2013, 400 pigs were sampled from seven sites across in Arua (Table 1).

**Table 1.**
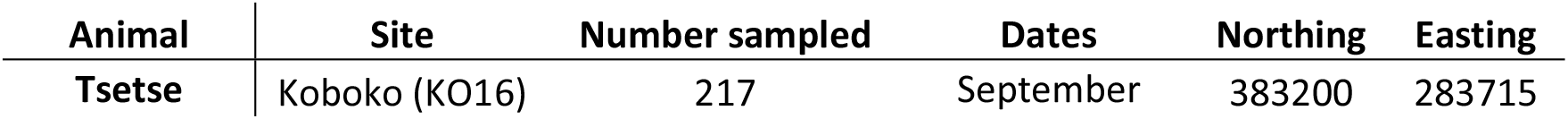
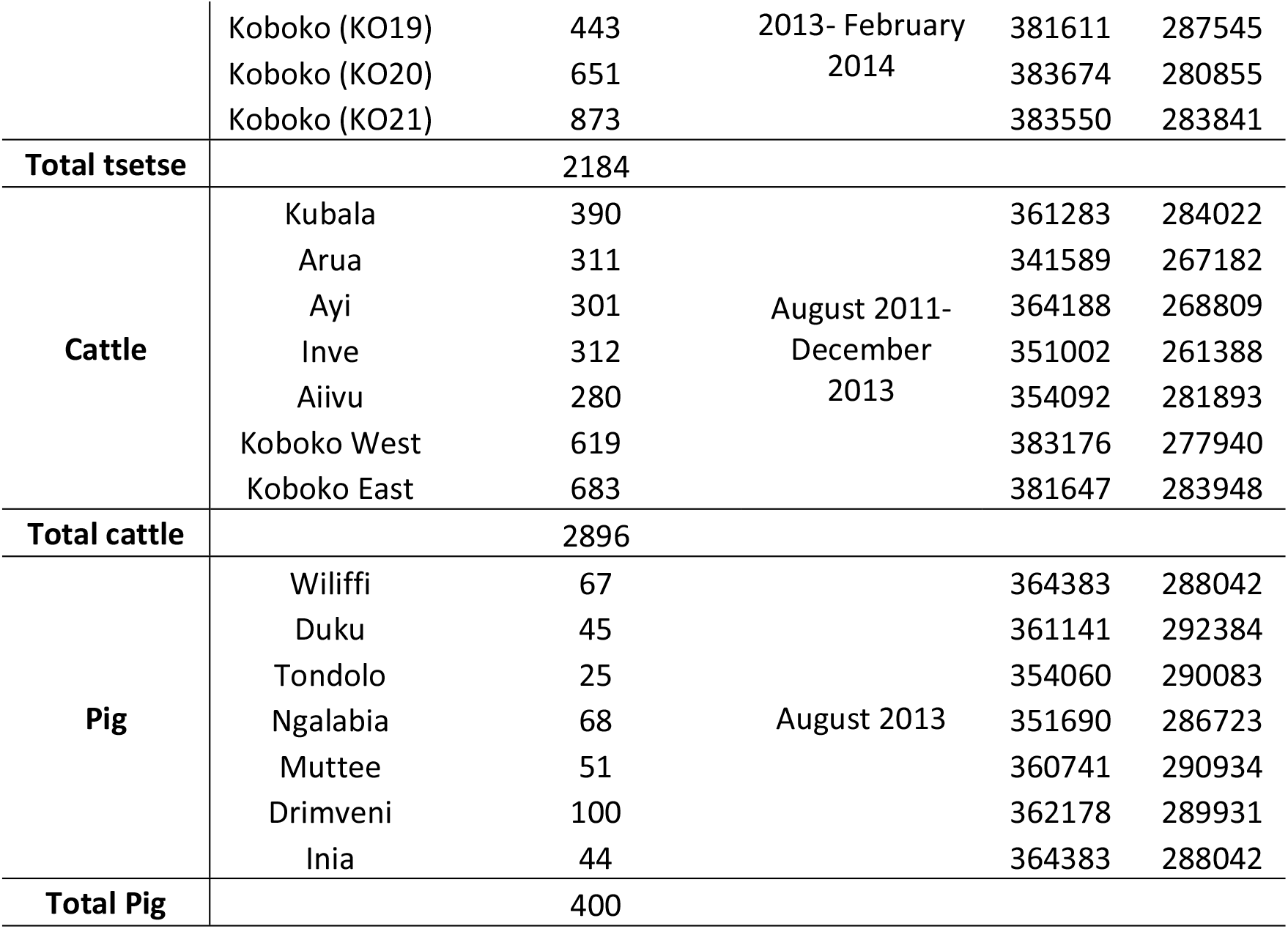
Tsetse, cattle and pig sampling sites with corresponding numbers of animals sampled

Both the cattle and pigs were sampled in the following manner, the animal was restrained, and a disposable lancet was used to puncture a pineal (ear) vein. Blood was collected with three 50mm heparinised capillary tube which collected 35μl of blood. Two tubes were centrifuged at 8,000 rpm for three minutes and the buffy coat layer examined as a wet preparation at x400 magnification using a compound-microscope with a dark-field filter. The contents of the third capillary tube was transferred to a Whatman FTA card (GE Health Care, Little Chalfont) and left to air dry before it was heat sealed in a foil pouch with a packet of silica gel to ensure the sample remained desiccated.

To extract the DNA from the FTA card, a modified version of the method described by Ahmed was carried out as follows, 10 2mm hole-punches were taken from each bloodspot, using a Harris micro-punch. The punches were washed three times in 1ml of distilled water and then 135μl of a 1% Proteinase K/10% Chelex TE suspension was added to each batch of 10-hole punches. These were then incubated at 56°C for an hour followed by 93°C for 30 minutes. In total 14 sampling sites (Fig. 1) were used to gather a total of 2896 samples from cattle and 400 from pigs across the N.W. of Uganda.

**Fig. 1.**
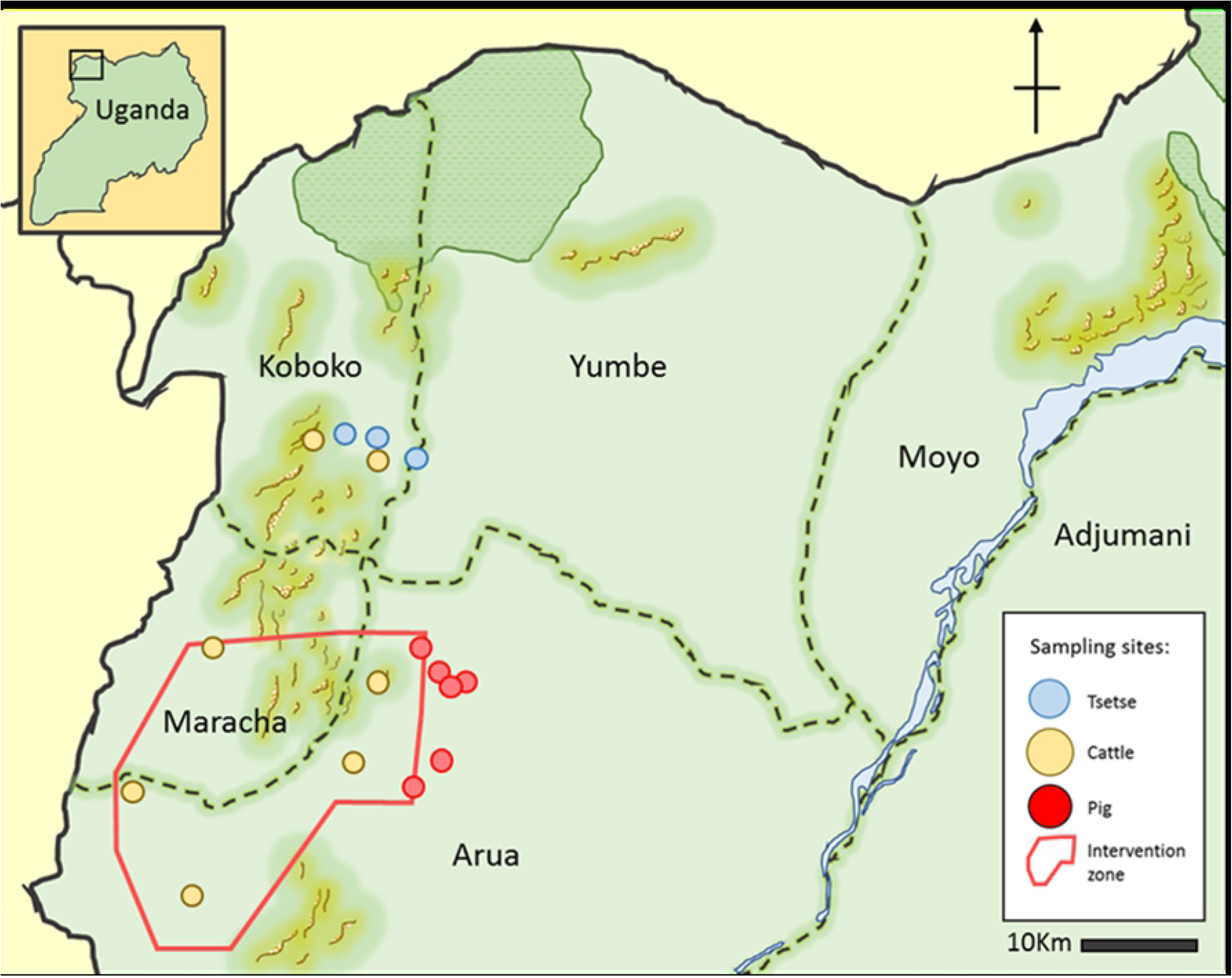
Map of sampling sites for tsetse, cattle and pigs from N.W. Uganda. Intervention zone denotes the area that was under tsetse control during the collection of samples described in this paper (27). The map was created by authors for this publication using Gnu Image Manipulation Software (31).

### Primer design

The tsetse and livestock samples underwent different PCR assays for the detection of *T. brucei* s.l.. Tsetse were processed with a nested multiplex primer set that targeted *T. brucei* s.l., *T. congolense* and *T. vivax* whereas cattle and pig samples were screened with the FIND-TBR primers (30). The nested multiplex primers were a modified version of the generic ITS primers designed by Adams *etal.* (32). These primers were used as part of a larger study to identify tsetse positive for *T. congolense* and *T. vivax* as well as *T. brucei*. The outer nest utilised the previously published forward and reverse primers that target the internal transcribed spacer region (ITS) of the trypanosome genome (32). New primers were designed to amplify species-specific regions from the first amplicon generated. This was achieved by aligning the ITS sequence for *T. congolense* Kilifi (accession number U22317), *T. congolense* Forest (accession number U22319), *T. congolense* Savannah (accession number U22315), *T. brucei* s.l. (accession number JX910373), *T. vivax* (accession number U22316), *T. godfreyi* (accession number JN673383) and *T. simiae* (accession number AB742533). A new universal forward primer and three species-specific reverse primers were designed and used in a multiplex. The location of the new primers are shown in a diagrammatic form in relation to the universal primers designed by Adams *eta*l. (Fig 2).

**Fig 2.**
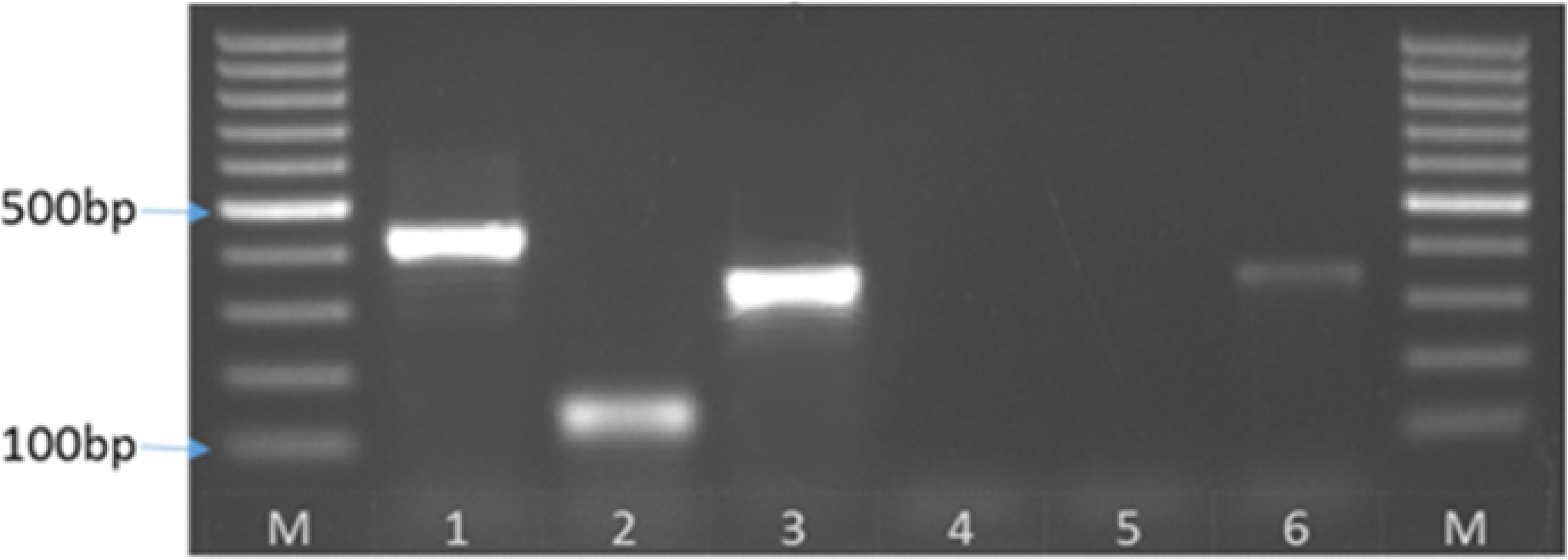
Diagrammatic representation of the location of the multiplex ITS primers in relation to the Universal primers designed by Adams et. al. (2006) on the ribosomal DNA.

The resulting products vary in size based on the species of trypanosome that was amplified with the largest product belonging to *T. congolense* s.p. measuring 392bp (*T. congolense* Kilifi) to 433bp (*T. congolense* Savannah and *T. congolense* Forest). The products for *T. brucei* s.l. and *T. vivax* measure 342bp and 139bp, respectively (Fig. 3).

**Fig. 3.**
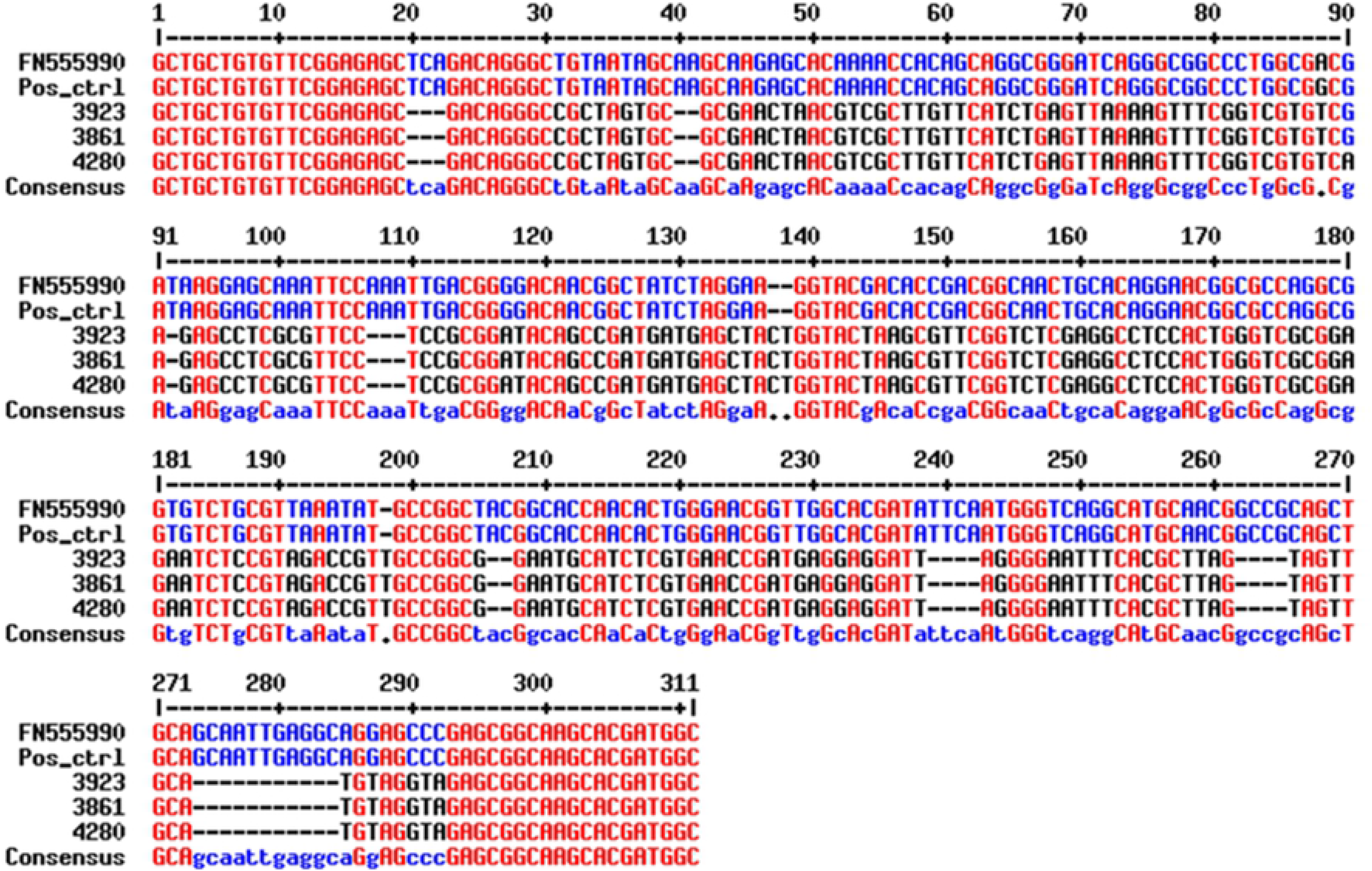
Image showing the relative sizes of the mITS PCR reaction for *T. congolense* Savannah (1), *T. vivax* (2) and *T. b. brucei* (3), extraction negative control (4), reagent negative control (5) and a second *T. b. brucei* positive control (6)

### *T. brucei* s.l. PCR reaction set-up

The PCR reactions were carried out at a final volume of 25μL each containing the following reagents: 2.5μl of 10X PCR Buffer (Bioline, London, UK), 200μM of each of the deoxynucleotide triphosphates (dNTPs) (Thermo Fisher Scientific, Leicstershire, UK), 1.2mM of MgCl2 (Bioline, London, UK), 0.4μM of both the forward and reverse primers and 10μL of BIOTAQ Red DNA Polymerase (Bioline). The first reaction of the two nested PCRs and standard PCR used 5μL of DNA template. For the nested PCRs second reaction 1μL of the PCR product from the first reaction was used as the template. The primers used to detect *T. brucei* s.l. positive samples are listed in Table 2. FIND-TBR primers were used for the cattle and pig samples and the novel multiplex ITS primers were used to screen the tsetse samples. The different strategies to screen for *T. brucei s.l.* reflects the different objectives of the animal and tsetse sampling with the focus on the animal samples being the identification of *T. brucei* positives while it was desirable to confirm presence of different species of Trypanosoma to support the tsetse microscopic examination of different tissues (midgut, mouthparts, salivary glands) from tsetse.

**Table 2.**
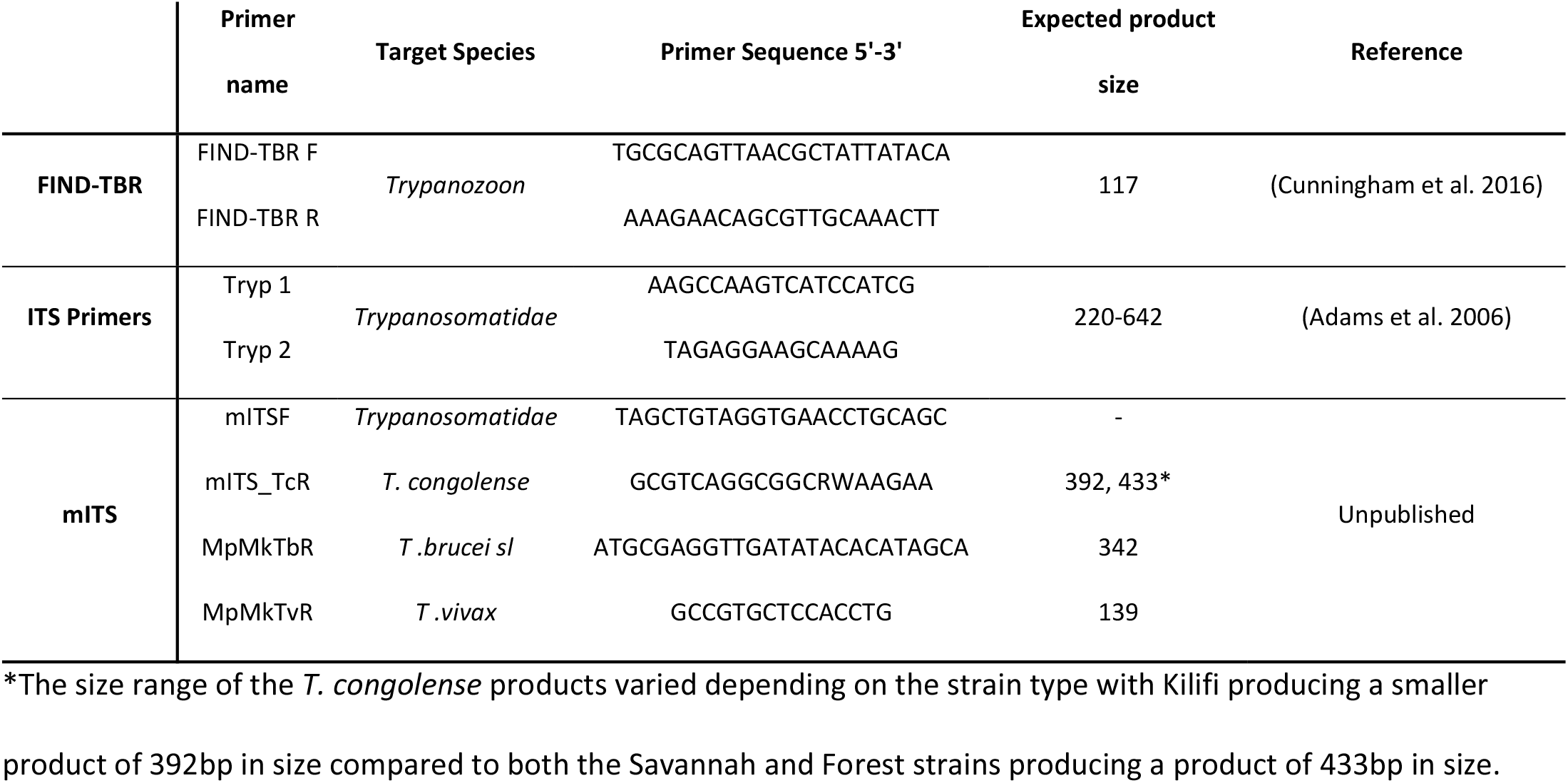
Primers used for detection of *T. brucei* s.l.

However as there is a difference in the copy number being targeted by the different primers not all those samples initially identified as *T. brucei s.l.* will be identified down to a sub-species level due to insufficient DNA. Following the detection of *T. brucei* s.l. positive samples using either the multiplex ITS primers or the FIND TBR-PCR primers (30), the positive samples were screened with the Picozzi multiplex primers (17). The FIND-TBR primer targets a copy region of several thousand (33) whereas the Picozzi primers and *T. b. gambiense* species specific primers (TgsGP) (34), target a single copy region of the genome. Therefore, it will not be possible to identify a proportion (^∼^27%) of the *T. brucei s.l.* positive samples down to the sub-species level as the species-specific primers are less sensitive than the TBR-PCR primers (35, 36). The following methodology was used to clarify how many TgsGP negatives were negative due to either the absence of *T. b. gambiense* or insufficient genomic material.

### Identification of single copy gene and *T. brucei* sub-species

Having identified which samples are positive for *T. brucei s.l.* there is a need to determine what sub-species of *T. brucei* the sample belongs, be it *T. b. brucei*, *T. b. gambiense* or *T. b. rhodesiense* due to the significance of the presence of a human infective sub-species. The PCR assays for positive identification of *T. b. rhodesiense* and *T. b. gambiense* rely on the amplification of a single copy gene unique to either the West or East African parasite. If neither Gambian or Rhodesian HAT is detected, then it is assumed the organism present is *T. b. brucei*. However due to the difficulty of reliably amplifying a single copy gene, and therefore the low sensitivity of the two sub-species specific assays, there is a chance that a negative result occurs due to insufficient target DNA. To help increase the confidence of a negative result it is possible to determine if there is sufficient quantity of DNA by running an assay that amplifies a single copy gene ubiquitous to all three sub-species. The multiplex designed by Picozzi *et al* (17) is capable of assessing whether there is sufficient DNA for single copy gene amplification and to also screen for *T. b. rhodesiense*. The multiplex consists of universal *Trypanozoon* primers that target the *T. brucei* s.l. single copy gene, phospholipase C (PLC), as well as primers that target the serum resistance associated gene (SRA) for *T. b. rhodesiense*. Among the variable surface glycoprotein (VSG) genes there are regions with some sequence identity to the *SRA* gene. Between the *VSG* and *SRA* genes there is an internal deletion within the *SRA* genes that allows it to be distinguished between any *VSG* amplification and *SRA* amplification. Two pairs of primers were designed that amplify across this deleted region to allow for clear size distinction between a *SRA* PCR product and a *VSG* PCR product. The combination of primers results in the amplification of a 324 bp product for all *Trypanozoon* species, a 669 bp product for *T. b. rhodesiense* and a >1 Kb for any *VSG* products amplified (Table 3).

**Table 3.**
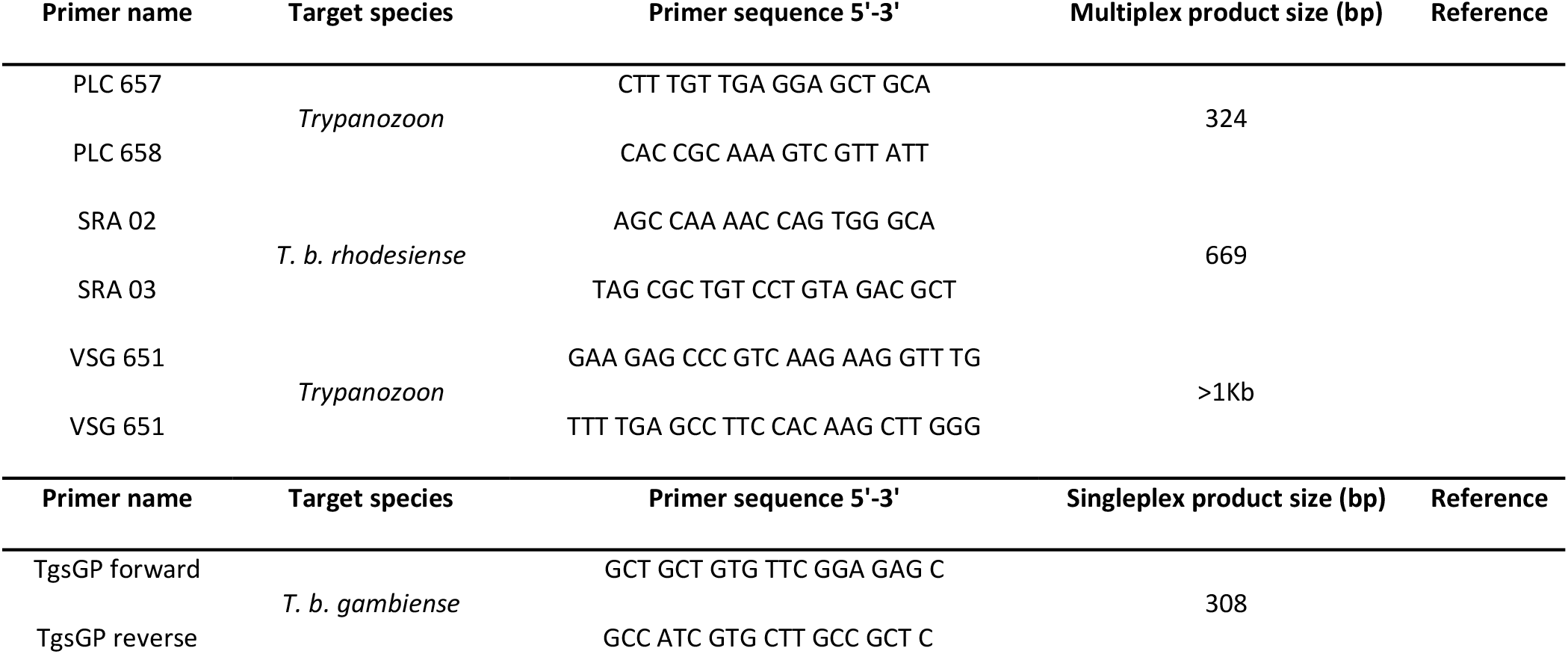
Primers for single copy gene detection and sub-species-specific analysis

### Ethics Statement

Ethics to sample domestic animals from Uganda was granted by the Ugandan National Council for Science and Technology ethics board, which approved the following protocol “Targeting tsetse: use of targets to eliminate African sleeping sickness” Ref. Number HS 939. The protocol followed the guidelines set out in the Ag Guide (37) and permission was granted by the animals owners for their involvement in the study.

## Results

### Microscopic examination

Of the 6,664 tsetse examined microscopically, 180 tsetse organs from 158 (2.4%; 95% CI=2-2.8) tsetse were positive for trypanosomes comprising 73 single midgut infections, 69 single mouthpart infections, nine mixed mouthpart-midgut infections, a single salivary gland-midgut infections and six cases where all three tissues were infected with trypanosomes. Of the 2,877 blood films examined from cattle, trypanosomes were identified in 568 (19.7%; 95% CI=18.3-21.2) samples, however, it was not possible to identify down to the species level using the microscopy methods.

### Molecular Screening for *T. brucei* s.l

In total 38/2,877 (1.3%; 95% CI=0.9-1.8) cattle and 25/400 (6.3%; 95% CI=4.1-9.1) pigs examined using the FIND-TBR primers were positive for *T. brucei s.l.*. The number of tsetse positive for the three Salivarian species of trypanosomes were as follows: *T. vivax* 46/2184 (2.1%; 95% CI=1.5-2.8), *T. brucei* s.l. 40/2184 (1.8%; 95% CI=1.3-2.5) and *T. congolense* 58/2184 (2.7%; 95% CI=2.0-3.4). Of the *T. brucei* positive tsetse, seven had a single positive mouthpart, nine were single salivary gland positives, 20 were single midgut positives, two were mixed salivary gland and midgut positive and the remaining two had all three tissues positive. The presence of *T. brucei s.l*. in the mouthparts deviates from the accepted life cycle of this trypanosome and is either due to a transient presence of trypanosomes that have passed along the proboscis during a feed or the result of cross contamination between tissues during dissection (Table 4).

**Table 4.**
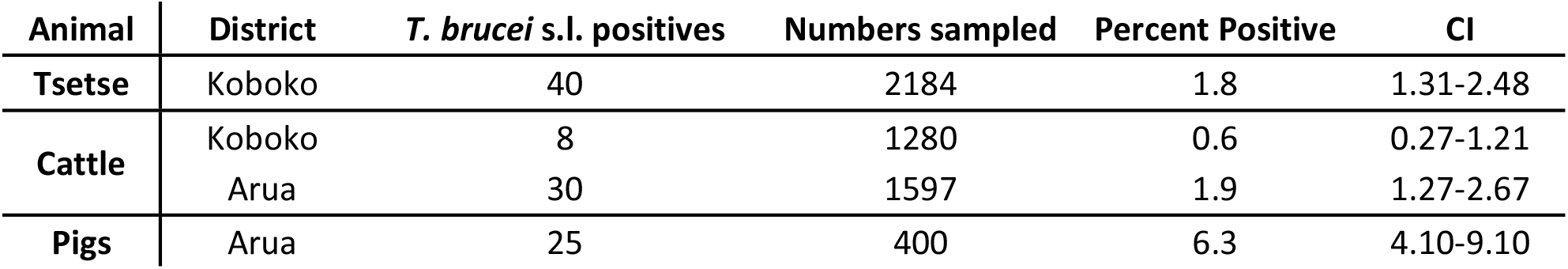
The prevalence of PCR *T. brucei* s.l. positives across all samples and districts.

| Animal | District | *T. brucei* s.l. positives | Numbers sampled | Percent Positive | CI |
| --- | --- | --- | --- | --- | --- |
| Tsetse | Koboko | 40 | 2184 | 1.8 | 1.31-2.48 |
| Cattle | Koboko | 8 | 1280 | 0.6 | 0.27-1.21 |
|  | Arua | 30 | 1597 | 1.9 | 1.27-2.67 |
| Pigs | Arua | 25 | 400 | 6.3 | 4.10-9.10 |

### Microscopy and Molecular comparison

Of the 2,184 samples screened with multiplex ITS primers, trypanosomes were observed by microscopy in 62 samples (49 flies), comprising 30 midguts, four salivary glands and 28 mouthparts. Of the microscopy positives the molecular assay identified 48 of 62 positive tissues, (Table 5).

**Table 5.**
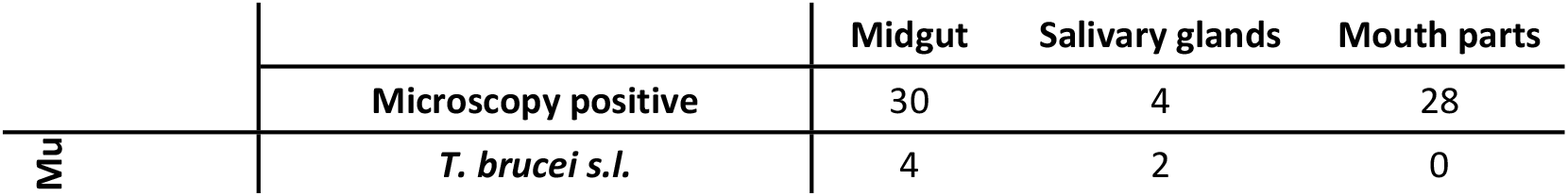
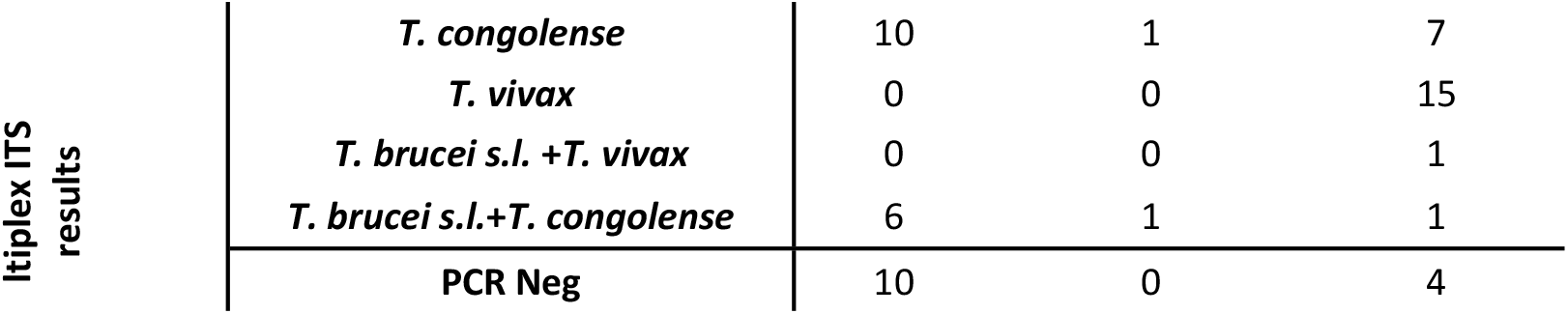
Species identification by multiplex ITS primers of microscopy positive tsetse tissues

### *T. brucei* s.l. sub-species screening

The 103 *T. brucei s.l*. positive cattle, pig and tsetse samples were processed using the *T. brucei* s.l. multiplex assay to screen for *T. b. gambiense*, *T. b. rhodesiense* and the number of samples with enough DNA to detect down to a single-copy gene. While not expecting to identify any cases of *T. b. rhodesiense* the inclusion of a positive control for the East African form of the disease acts as another quality control (Fig 4). None of the samples tested positive for the SRA gene, confirming our expectation that T. b. rhodesiense was not present. However of the cattle, pigs and tsetse, 24 (63%), 15 (60%) and 25 (56%) were positive for the PLC gene, respectively, suggesting that sufficient DNA was present to detect DNA from T. b. rhodesiense or *T. b. gambiense* if it were present.

**Fig 4.**
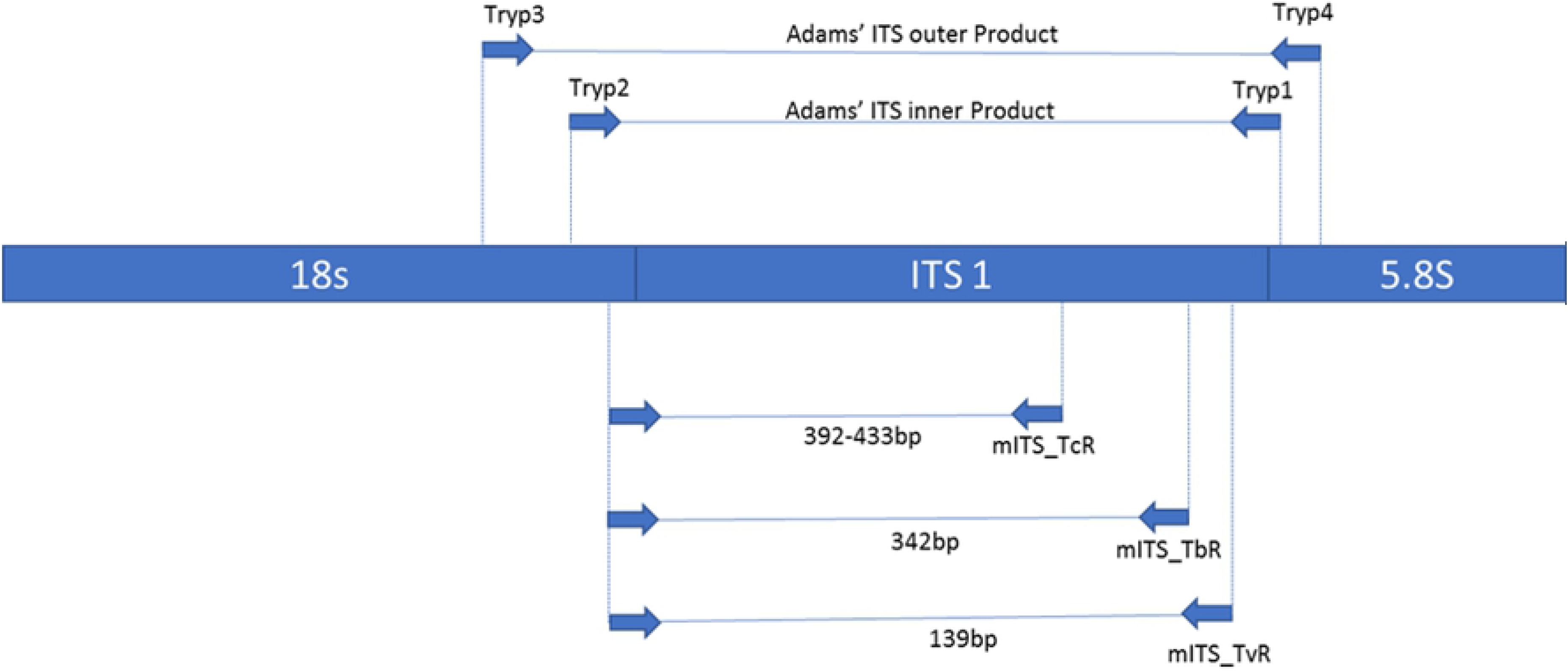
Example of gel run showing the results of the sub-species *Trypanozoon* multiplex PCR. The 324bp PLC product can be seen in the three positive controls and in sample number 2. The 669bp SRA product can be seen in the *T. b. rhodesiense* positive control but is absent from the other *T. brucei* sub-species. A >1 kb VSG product is visible in sample 2 and is faintly visible in the *T. b. rhodesiense* positive control. A fourth product of ^∼^700 bp can also be seen in the *T. b. gambiense* and *T. b. rhodesiense* positive controls as well as sample 2.

The samples that proved to have enough genetic material for the amplification of the single copy PLC gene were then screened using the sub-species-specific *T. b. gambiense* primer, TgsGP. Among the cattle and pig samples there was no amplification of a 308 bp product however 17 tsetse samples produced a band approximately 308 bp in size.

Of the 17 bands, a subsample of three were sent for sequencing to determine the specific product size and sequence. The samples were sent to SourceBioscience using both forward and reverse primers. The results of the sequencing showed that the three bands sent were identical and that the product was 281 bp (inclusive of primers) in length. The sequence when aligned against reference sequences for *T. b. gambiense* using the NCBI database resulted in only a 90% identity and a query cover of 16% Fig 5.

**Fig 5.**
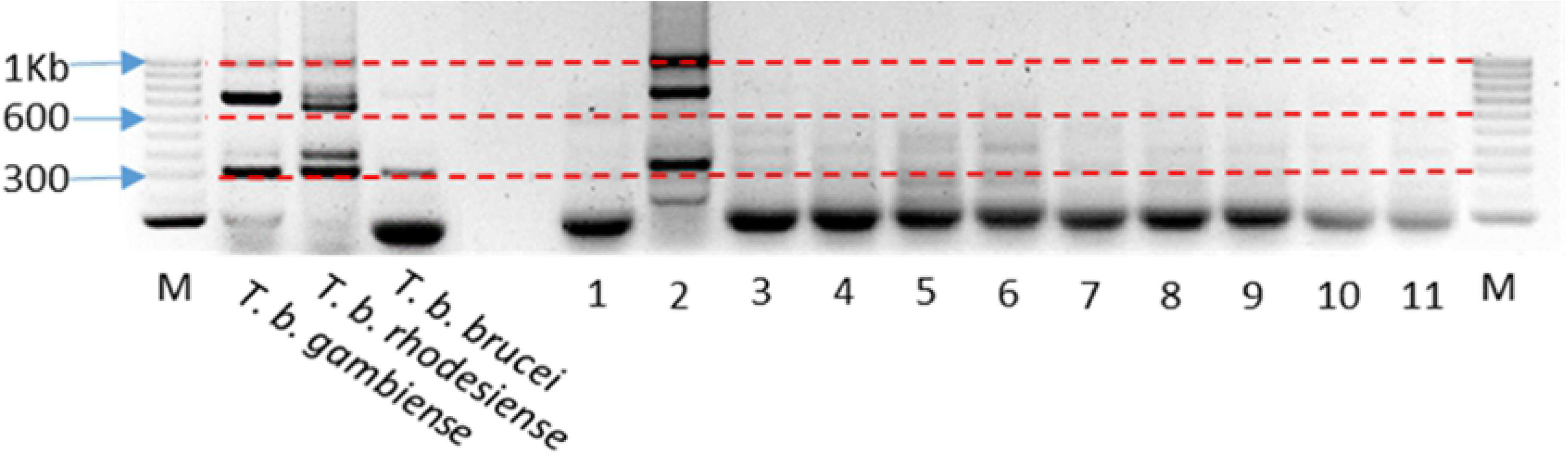
Results of the sequence alignment of the three samples sent for sequencing (3923, 3861 and 4280) against the *T. b. gambiense* positive control (Pos_ctrl) and reference sequence (FN555990) from the NCBI database. Image generated using MultAlin (38).

## Discussion

The aim of this paper was to determine the prevalence of *T. b. gambiense* in local tsetse, cattle and pig populations from N.W. Uganda. The successful identification of *T. b. gambiense* would suggest that transmission of sleeping sickness in the area was continuing and the identification of the disease in either cattle or pigs would help resolve the role of animal reservoirs in the transmission of the disease.

### Tsetse

Of the 40 tsetse samples identified as *T. brucei* s.l. positive by PCR 56% were found to have enough DNA for the amplification of a single-copy gene region. Sixteen produced faint bands of approximately 300bp, comparable in size to the expected band size for *T. b. gambiense*. However sequencing results showed that the size of the product generated by the samples was smaller than the expected size at only 281bp compared to the expected 308bp sized product. There was also significant variation in the 281bp sized sequences compared to *T. b. gambiense* sequence. Based on the sequencing results these positive samples cannot be unequivocally identified as *T. b. gambiense*.

Three conclusions arise from the tsetse survey, the first is that despite screening 2,184 tsetse, no tsetse were found to be positive for *T. brucei gambiense*, this could be due to *T. b. gambiense* no longer being transmitted in the area or that our sample size was too small to detect *T. b. gambiense*. Despite being understood as the sole vector of gHAT (39) the prevalence of the disease amongst wild tsetse population is often extremely low (1, 40, 41) and attempts to infect tsetse with *T. b. gambiense* under laboratory conditions have often proven unsuccessful (42). Studies have suggested that the prevalence of *T. b. gambiense* may be as low as 1 in 4,000 flies (11). However, this number is based on microscopy methods, whereas PCR methods should be more sensitive and could identify immature and transient infections reducing the number of tsetse needed to be screened (8). Second despite no *T. b. gambiense* being found, the tsetse population studied were actively transmitting *T. b. brucei*. Third, the TgsGP primers cross-reacted with DNA from an unidentified source and produced a band, similar in size, to *T. b. gambiense*, this raises concerns about the specificity of the TgsGP primers are and the potential for erroneously reporting the presence of *T. b. gambiense*

Table 4 shows that the positive midguts were identified as either *T. brucei* or *T. congolense* similarly of the positive mouthparts all were infected with either *T. congolense* or *T. vivax* with no mono infections of *T. brucei* s.l. identified, although in cases of mixed infections *T. brucei* s.l. was detected in the mouthparts.

The infected salivary glands were predominantly positive for *T. brucei* s.l. however there was one instance of a single *T. congolense* infection. The presence of a mITS positive does not indicate an infection of a specific tissue by the trypanosome detected but merely the DNA, which could be a transient trypanosome, free floating DNA or DNA introduced during the dissection step; previous studies have found similar results (43). Although, overall, the comparison between the mITS results and those of the dissection correlate closely with the traditional methods used to identify trypanosome species based on their location in different tsetse tissues (29), however these methods cannot distinguish between species easily and certainly not between sub-species.

### Cattle and pigs

No animal samples (pig or cattle) produced a band of either 308bp or 281bp when screened with the TgsGP primers, indicating that there were no zoonotic *T. b. gambiense* infections nor where there any cases similar to those found in the fly samples, where non-target DNA was amplified. However, cases of *T. b. brucei* were found in both animal populations with pigs proving to have the higher prevalence of *T. b. brucei* infection. This is typical of trypanosome epidemiology which has been shown to be highly localised in other studies (44). The lack of any positive *T. b. gambiense* in the two animal populations sampled correlates with a previous study carried out in the same area (18), indicating that it is unlikely either cattle or pigs are acting as cryptic reservoirs of disease.

### Diagnostics for *T. b. gambiense*

The diagnostic methods used in this paper involved both microscopy and PCR, of which only PCR has the potential to discriminate sub-species of *T. b. gambiense* (17, 34). There are few diagnostic methods that are capable of accurately distinguishing between the *T. brucei* sub-species (16, 17, 34). The molecular methods available for the detection of *T. b. gambiense* are limited due to the practical aspect of conducting these assays in the field and the limited diagnostic markers available. As mentioned previously the sensitivity of *T. b. gambiense* specific PCR is limited to detecting a single copy gene. Some molecular assays attempt to overcome this problem by relying on the human serums ability to lyse all salivarian trypanosomes (except for *T. b. gambiense* and *T. b. rhodesiense*) therefore any *T. brucei* s.l. identified in a human sample would be one of the two HAT species (45). This allows for the targeting of a higher copy region specific to the *T. brucei* species group. Using this method any positives would have to be one of the two HAT species, however as the treatment of the two diseases differs and the only option to try and identify if it was an East or West African sleeping sickness infection would be to try and determine the geographical location of where the individual was infected. This approach would also only be limited to humans and could not be used in either xenodiagnoses or screening animals, as all three *T. brucei* sub-species could be present in the vector or animal populations. To put these differences of sensitivity in perspective, we can look at the limit of detection (LoD) of the number of trypanosome per mL, between multiple diagnostic methods (Table 6.).

**Table 6.**
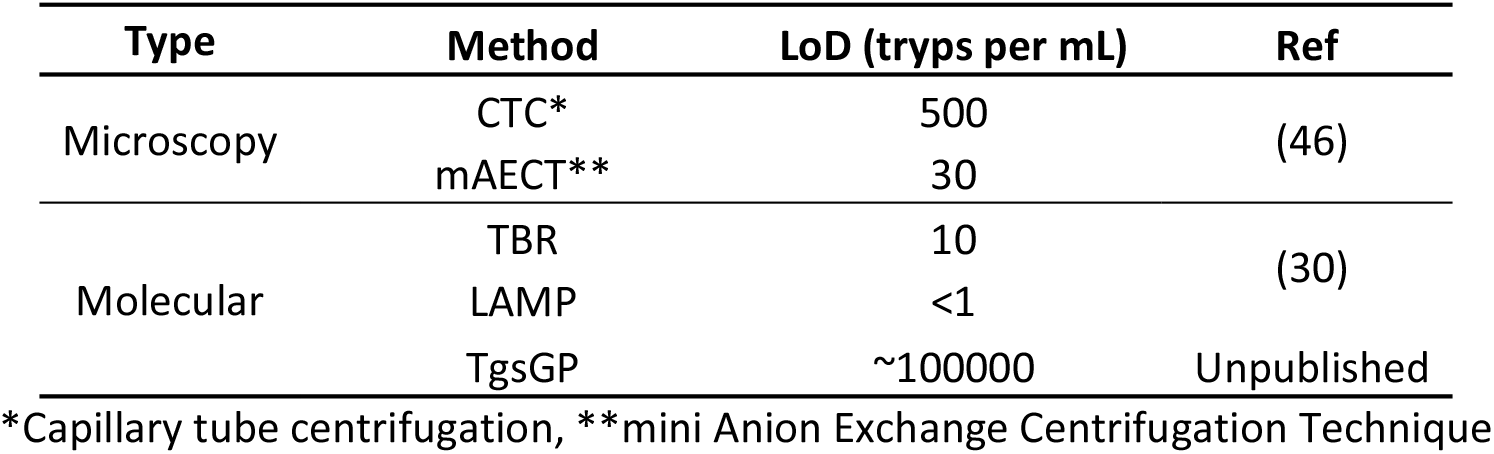
Comparison of LoD between different diagnostic techniques

The lack of a highly specific, sensitive and field-friendly assay that is capable of screening for *T. b. gambiense* in both the human, vector and local animal populations is sorely needed if the hope of eliminating sleeping sickness by 2020 is to be achieved.

## Conclusion

This lack of positive samples reflects the overall low prevalence of the disease and the continued decrease in the number of cases in Uganda (47). This study also highlights the lack of highly sensitive diagnostics that can discriminate between the different sub-species of *T. brucei* s.l.. Despite not finding *T. b. gambiense* in the tsetse population of Koboko vector control has been calculated to being essential to reach the elimination goal of 2030 (48)

## Acknowledgments

The authors of this paper would like to thank the staff based in Uganda and LSTM that assisted in the fieldwork, notably Victor Drapari, Edward Aziku, Henry Ombanya.

